# Non-invasive tracking of ripple-like activity during human sleep using MEG

**DOI:** 10.64898/2026.07.20.739563

**Authors:** Fabian Schwimmbeck, Jan Martini, Christian Vollmar, Tobias Staudigl, Eugen Trinka, Jan Rémi, Christian F. Doeller, Ole Jensen, Thomas Schreiner

## Abstract

Hippocampal ripples are considered a key mechanism of sleep-dependent memory consolidation. In humans, however, their direct investigation has relied on invasive recordings from patients with epilepsy, because EEG provides limited access to fast activity from deep medial temporal structures. Here, we tested whether source-resolved magnetoencephalography can reveal physiologically meaningful ripple-like activity during NREM sleep in healthy humans. These events showed hallmarks expected of sleep ripples: they were most prominent in medial temporal regions and embedded within slow oscillation (SO)–spindle dynamics, occurring preferentially during SO up-states and spindle centers. To test mnemonic relevance, we combined sleep MEG with targeted memory reactivation, in which learning cues were replayed during sleep. TMR improved memory performance, and later remembered cues preferentially recruited medial temporal ripple-like events alongside cortical spindles. Ripple-like events also coincided with enhanced item-specific reactivation during successful TMR. Together, these findings establish MEG as a non-invasive window onto hippocampo-cortical memory dynamics during sleep.

## INTRODUCTION

Non-rapid eye movement (NREM) sleep is thought to support the consolidation of newly acquired memories through the reactivation of information encoded during prior wakefulness^1–3^. Through repeated reactivation, initially labile hippocampus-dependent memories are gradually stabilized and integrated into neocortical networks^4–7^. This process is coordinated by a canonical oscillatory hierarchy comprising cortical slow oscillations (SOs, <1 Hz), thalamo-cortical sleep spindles (12-16 Hz), and hippocampal sharp-wave ripples (80-120 Hz in humans^8–11^. In this context, hippocampal ripples are closely linked to the reactivation of recent experiences and are widely considered a central mechanism of sleep-dependent memory consolidation^12–16^. Their temporal coordination with SOs and spindles is thought to promote hippocampo-cortical communication and synaptic plasticity, thereby supporting the stabilization of memory representations in distributed cortical networks^17–19^.

This mechanistic account is strongly supported by rodent work, in which disruption of hippocampal sharp-wave ripples impairs memory consolidation and coordinated SO–spindle-ripple coupling predicts successful systems-level memory processing^7,14,17,20^. In humans, however, direct investigation of hippocampal ripple dynamics has remained largely limited to intracranial recordings in epilepsy patients undergoing presurgical monitoring^21^. Although scalp EEG is the gold standard for sleep research and ideally suited for tracking cortical sleep oscillations^22^, it provides only limited access to the fast ripple-frequency activity generated in deep medial temporal structures, because these signals are strongly attenuated and spatially blurred by volume conduction through the skull and surrounding tissues^23^. Intracranial studies in patients with epilepsy have nevertheless provided converging evidence in humans, showing that hippocampal ripples are embedded within the canonical SO-spindle architecture of NREM sleep^8,11,16,24,25^ and that this coordinated interplay is linked to memory reactivation and sleep-dependent consolidation^15,16^.

At the same time, although invasive studies have provided critical insights into human ripple physiology, the clinical setting imposes important limitations on establishing how such dynamics operate in the healthy human brain^21^. Electrode coverage is determined by clinical rather than experimental considerations. As a result, spatial sampling is sparse and heterogeneous across patients. Moreover, ripple-frequency activity may be influenced by underlying pathology or intermixed with epileptiform high-frequency events, complicating the separation of physiological ripples from pathological activity^26–29^. As a result, it remains unclear to what extent the canonical SO-spindle-ripple architecture established in animal models and intracranial human studies generalizes to healthy human sleep.

Crucially, although hippocampal signals were long considered difficult to detect non-invasively because of their depth and presumed closed-field geometry, a growing body of computational and empirical work suggests that MEG , particularly when combined with source-level analyses, can provide access to neural activity arising from deep medial temporal lobe (MTL) structures, including the hippocampus^30–32^. Compared with EEG, MEG may offer a particular advantage because magnetic fields are less distorted by the skull and surrounding tissue boundaries, potentially preserving the temporal and spatial characteristics of fast neural activity more faithfully at the sensor level^23^. Consistent with this idea, a growing number of source-resolved MEG studies have linked hippocampal or broader MTL activity to episodic memory, working memory, and spatial navigation tasks^30,33–44^. Together, these findings suggest that source-level MEG may capture functionally meaningful MTL activity in humans and raise the possibility that it may also provide access to ripple-like activity during NREM sleep.

A key challenge, however, is to establish whether fast events detected in source-reconstructed MEG data reflect physiologically meaningful ripple-like activity. Sleep offers a particularly powerful validation context, because hippocampal ripples have well-characterized spatial, temporal, and functional signatures from animal electrophysiology and human intracranial recordings^8,11,12,15,16,20,45^. Ripple-like MEG activity should therefore converge with these established signatures: it should arise from medial temporal regions^2,12,46–48^, be embedded within the canonical SO–spindle architecture of NREM sleep^10,11^, and relate to memory processing^12–14,17^. Such convergence would support the physiological relevance of source-reconstructed fast MEG events, positioning MEG as a promising non-invasive window onto ripple-like medial temporal dynamics during human sleep.

Building on this validation framework, we tested whether MEG-detected events during human NREM sleep show the spatial, temporal, and functional profile expected of hippocampal ripples. We identified ripple-like events in medial temporal source space and examined their temporal coupling to cortical SOs and sleep spindles. To probe their functional relevance for sleep-dependent memory processing we additionally leveraged targeted memory reactivation (TMR), an approach in which auditory memory cues are re-presented during NREM sleep to experimentally induce memory reactivation^15,49–54^. We found that endogenous ripple-like MEG activity was preferentially localized to medial temporal regions and recapitulated the established temporal relationship to canonical sleep oscillations, occurring preferentially around SO-upstates and sleep spindles during sleep. Importantly, the coupling dynamics of the source-localized MEG ripples matched those observed in a complementary intracranial EEG (iEEG) analysis. Moreover, in the TMR condition, ripple-like activity was selectively recruited together with cortical spindles during successful memory cueing. These findings suggest that MEG provides a non-invasive window onto ripple-like medial temporal dynamics in healthy humans and offers a scalable approach for studying hippocampo-cortical coordination during sleep-dependent memory consolidation.

## RESULTS

We combined simultaneous MEG and EEG sleep recordings with a TMR paradigm to examine whether MEG can reveal physiologically meaningful ripple-like activity during human NREM sleep. Following our validation framework, we assessed whether MEG-detected ripple-like events exhibited three established signatures of hippocampal ripples: (i) medial temporal localization, (ii) embedding within canonical NREM sleep oscillations, and (iii) functional relevance for memory processing. Eighteen healthy, right-handed participants took part in the study (age: 23.94 ± 0.97 years, mean ± SEM; 9 female). They completed two experimental sessions separated by approximately one week: one non-TMR session without cue presentation (n = 18) and one TMR session, in which auditory memory cues were presented during sleep (n = 16 after exclusion of two sessions with insufficient usable TMR data). In both sessions, an episodic memory task was performed before a 90-min nighttime nap in the MEG (for sleep characteristics see Supplementary Table 1), with memory performance assessed immediately before sleep (Recall Pre) and after sleep (Recall Post; Fig. 1A; see Supplementary Table 1 for sleep parameters). During encoding, participants learned associations between spoken verbs and visually presented objects. In the TMR session, auditory cues corresponding to half of the previously learned items (70 out of 140) were replayed during NREM sleep, with each item presented six to seven times (mean ± SEM = 6.88 ± 0.35). No reminder cues were presented during the non-TMR session.

**Figure 1.**
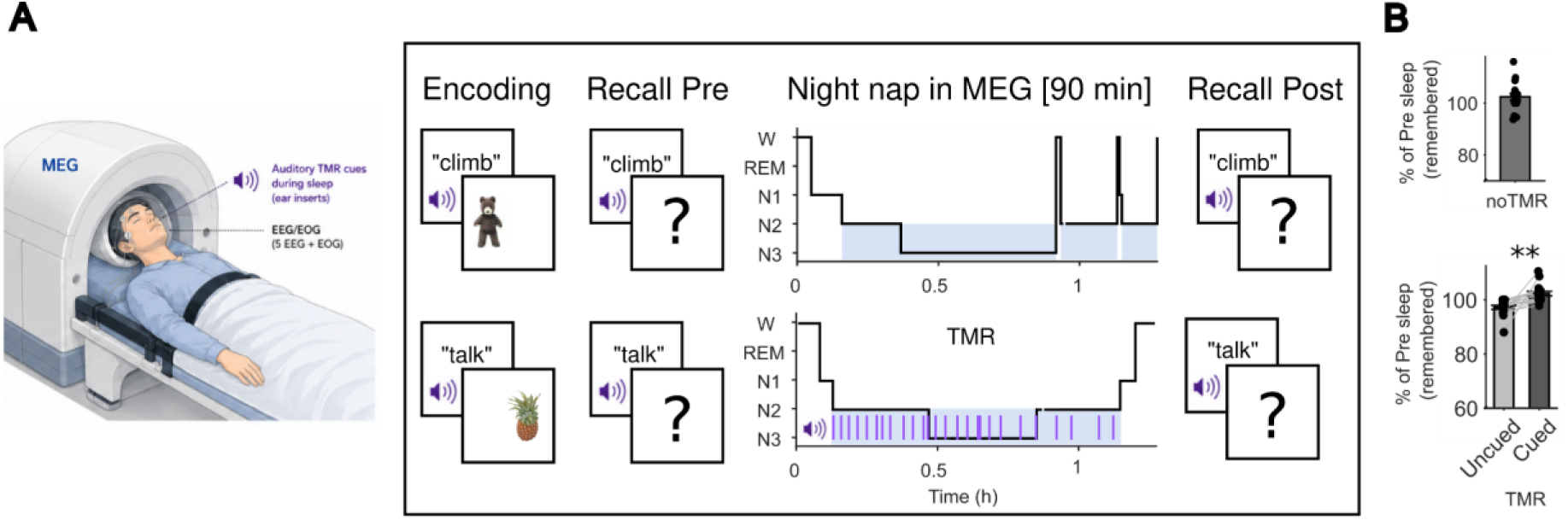
Experimental procedure and behavioural outcome. **A**, Experimental design. Left, schematic illustration of the simultaneous MEG-EEG sleep-recording setup used during the 90-min nighttime nap. Middle, task information and experimental timeline. Participants completed two sessions separated by approximately one week: a TMR session and a non-TMR control session (counterbalanced order). Before sleep, participants learned associations between spoken verbs and visually presented objects and completed an initial memory test (Recall Pre). In the TMR session, auditory cues corresponding to half of the learned associations were repeatedly presented during NREM sleep, whereas no cues were presented in the non-TMR session. Memory was reassessed after sleep (Recall Post). **B,** Behavioural memory performance. Upper panel, memory retention in the non-TMR session expressed as post-sleep recall relative to pre-sleep recall (Post/Pre × 100). Memory performance was preserved across the nighttime nap and did not differ significantly from 100% retention (two-sided one-sample *t*-test, *p* = 0.074, n = 18). Lower panel, memory retention in the TMR session for cued and uncued associations. TMR benefited subsequent memory retention: items cued during sleep were remembered better than uncued items (two-sided paired *t*-test, *p* = 0.0018, n = 16), demonstrating a behavioural benefit of TMR. Data are shown as individual participants with group mean ± SEM. Figure A was created using ChatGPT (OpenAI) and edited by the authors.

### Behavioural Results

We first examined whether memory performance changed across the nighttime nap and whether TMR influenced subsequent memory retention. In the non-TMR session, overall memory retention was quantified as post-sleep recall relative to pre-sleep recall (Post/Pre × 100), with 100% reflecting unchanged recall across sleep. Recall was largely preserved across sleep, with post-sleep performance remaining close to pre-sleep levels (mean ± SEM = 102.44 ± 1.28%) and not differing significantly from 100% retention (two-sided one-sample t-test: t(17) = 1.91, *p* = 0.074; Fig. 1B, upper panel). In the TMR session, overnight retention differed significantly between cued and uncued items. Recall was higher for cued than uncued associations (cued: mean ± SEM = 102.32 ± 0.84%; uncued: mean ± SEM = 97.21 ± 0.75%; mean difference ± SEM = 5.10 ± 1.35%; two-sided paired t-test: t(15) = 3.77, *p* = 0.0018, n = 16; Fig. 1B, lower panel), indicating a behavioural benefit of TMR. This cueing-related memory advantage provided the behavioural basis for later analyses linking successful cueing to sleep oscillatory dynamics and ripple-like MEG activity.

### MEG resolves canonical sleep oscillations and their coupling

We first examined whether MEG captured the canonical oscillatory structure of NREM sleep, providing the physiological context for subsequent analyses of ripple-like activity. To visualize spectral dynamics during NREM sleep, we applied multi-taper spectral analyses to EEG and MEG signals. Across recording modalities and analysis levels, spectral sleep profiles were preserved from scalp EEG (C4) to MEG sensor space (frontal sensor MLF12) and hippocampal source space, with all modalities exhibiting canonical slow-wave (∼1 Hz) and spindle-band (∼15 Hz) activity characteristic of NREM sleep (Fig. 2A, example participant). This preservation of NREM spectral signatures indicates that sleep-related oscillatory dynamics remain detectable even after MEG source reconstruction into deep brain regions. We next detected cortical slow oscillations (SOs; mean ± SEM = 398.62 ± 30.55 events per subject and night nap, 2.86 ± 0.14 events/30 s; Fig. 2B) and sleep spindles (mean ± SEM = 578.19 ± 32.42 events per subject and night nap; 4.20 ± 0.14 events/30 s; Fig. 2C) in MEG sensor space using established detection algorithms ^11,25,55^, with the corresponding sensor-level topographical distributions shown in Supplementary Fig. 1 SOs and spindles showed the expected temporal coupling, with spindle activity preferentially nesting within SO up-states (*p* = 0.025, −0.80 to −0.40 s relative to SO-trough, and *p* = 0.002, 0 to 0.55 s relative to SO-trough, corrected across time and sensors; Fig. 2D, left). These coupling effects were most prominent over frontal, temporal, and parietal-occipital sensor regions (Fig. 2D, middle). Complementing these time-resolved analyses, a phase-domain analysis was used to determine the preferred SO phase of spindle occurrence. Spindle occurrence was non-uniformly distributed across the SO cycle and significantly aligned to the SO up-state (V-test against 0°, where 0 corresponds to the SO up-state: mean angle ± SEM = 17.31° ± 6.65°, v = 15.10, *p* < 0.0001, ; Fig. 2D, right). Together, these findings establish that sleep MEG reliably captures canonical NREM sleep oscillations and their characteristic SO–spindle coupling.

**Figure 2.**
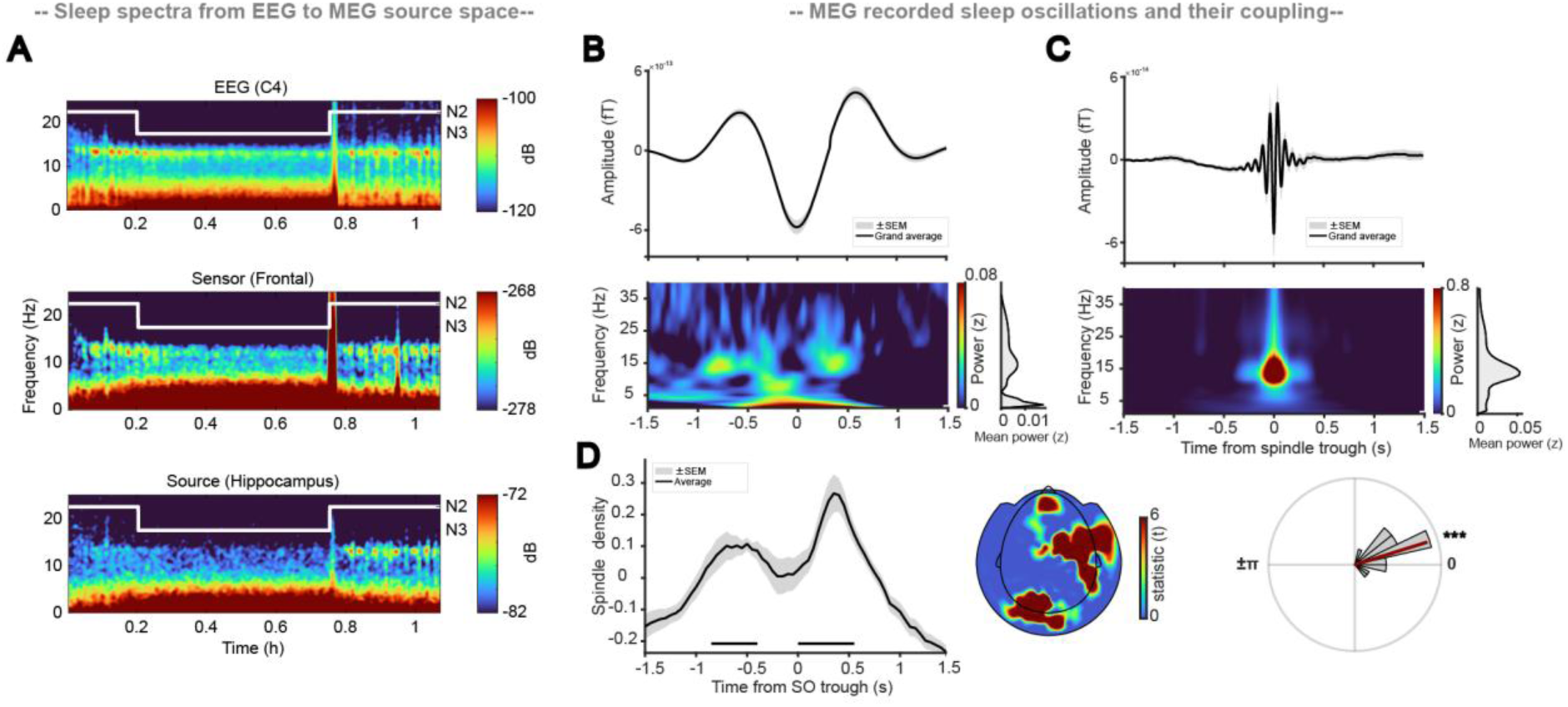
MEG resolves canonical NREM sleep oscillations and their coupling. **A**, Spectral profiles across recording modalities and analysis levels, from scalp EEG (central electrode C4) to MEG sensor space (frontal sensor MLF12) and MEG source space (hippocampus) during NREM sleep in a sample participant. Spectrograms consistently revealed canonical slow-wave activity (∼1 Hz) and spindle-band activity (∼15 Hz) across modalities, demonstrating preservation of characteristic NREM oscillations following MEG source reconstruction. **B,** MEG sensor-level SOs. Top, grand-average SO waveform time-locked to the SO trough across sessions. Bottom, SO-trough-locked TFR. A total of 398.62 ± 30.55 SOs (2.86 ± 0.14 events/30 s) were detected per participant (mean ± SEM.). **C,** MEG sensor-level spindles. Top, grand-average spindle waveform time-locked to the spindle trough, defined as the most negative deflection, across sessions. Bottom, spindle-trough-locked time–frequency representation. A total of 578.19 ± 32.42 spindles (4.20 ± 0.14 events/30 s) were detected per participant (mean ± SEM.). **D,** Coupling between SOs and sleep spindles. Left, spindle occurrence relative to SO troughs, showing preferential spindle recruitment during the first and second SO up-state (*p* = 0.025, −0.80 to −0.40 s, and *p* = 0.002, 0 to 0.55 s; corrected across time and sensors). Middle, topographical distribution of significant SO-spindle coupling effects, maximal over frontal, temporal and parietal-occipital sensors. Right, distribution of spindle occurrence across the SO phase cycle, demonstrating significant alignment to the SO up-state (V-test against 0°, where 0 corresponds to the SO up-state: mean angle ± SEM = 17.31° ± 6.65°, *v* = 15.10, *p* < 0.0001).

### NREM MEG ripple-like activity localizes to the MTL and couples to canonical sleep oscillations

Genuine ripple activity would be expected to arise from medial temporal regions and to be temporally coordinated with cortical SOs and sleep spindles, as shown in rodent electrophysiology and human intracranial EEG recordings^7,8,10,11,14–16^.

To test this, we reconstructed voxel-wise source time series from the MEG sensor data using an LCMV beamformer^56^ and applied ripple-detection procedures during NREM sleep, using criteria analogous to established intracranial EEG approaches^8,11,15,16^. Ripple-like events were first detected independently across a whole-brain source grid comprising 1,457 voxels. For the initial source-space time–frequency analysis, we derived ripple-associated episodes from neighbouring voxel-level detections, providing a common event set for comparing ripple-band power across source voxels and identifying spatial hotspots of ripple-like activity (see Methods). Compared with surrogate events, these ripple-associated episodes showed a spatially selective increase in high-frequency power, with significant clusters peaking at 94 Hz and centered on the MTL and downstream temporal regions (p = 0.002, 10.99 to 150.02 Hz, −0.200 to 0.500 s from ripple center, corrected across time, frequency, and voxels). To focus subsequent analyses on voxels with the clearest ripple-like signal, we defined a data-driven hotspot mask comprising the 15% of voxels with the strongest ripple-band power increases relative to surrogate events (n = 219; Fig. 3A, bottom; the full unmasked source-space statistical map is shown in Supplementary Fig. 2A). Within these hotspot voxels, ripple-like events showed pronounced transient increases in 80–120 Hz power (Fig. 3A, top and middle), yielding 193.71 ± 24.28 events (mean ± SEM, 1.39 ± 0.16 events/30 s). An independent ripple-density-based analysis on all voxels (mean ± SEM = 1.19 ± 0.11 events/30 s) revealed a convergent spatial distribution, with peak event density likewise observed in the MTL and downstream temporal regions (Supplementary Fig. 2B). Thus, converging power- and event-based analyses support the spatial plausibility of MEG ripple-like activity as reflecting medial temporal sources.

**Figure 3.**
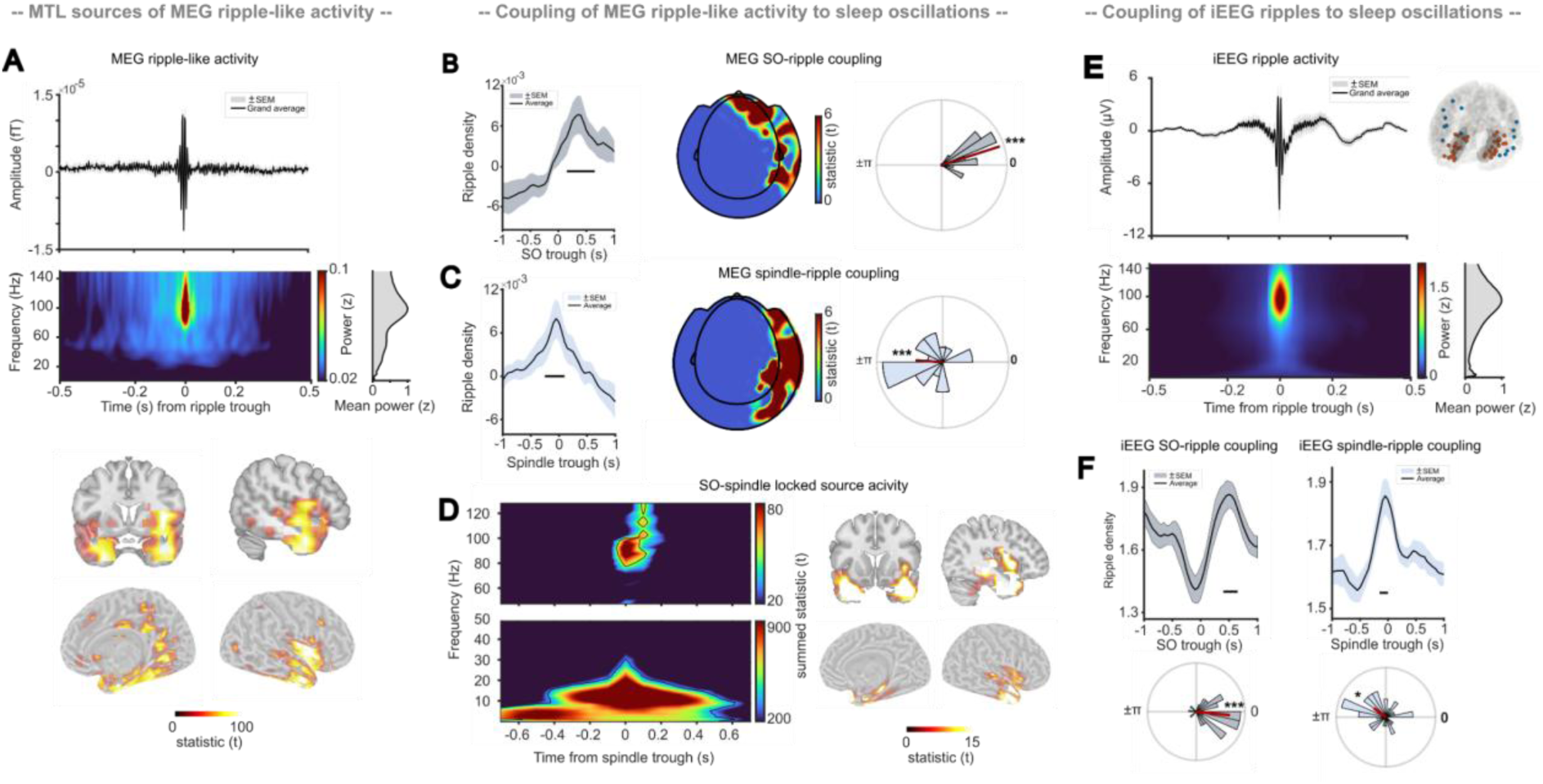
MEG ripple-like activity shows MTL origin and coupling to cortical oscillations. **A**, Grand-average ripple-like activity detected voxelwise in MEG source space time series. Top, ripple-triggered ERP within hotspot voxels (n=219, see below). Middle, TFR of ripple-like events. Bottom, cluster-based permutation analysis identifying significant ripple-band power increases ) during ripple episodes, with effects peaking at 94 Hz relative to surrogates, localized primarily to MTL and downstream temporal regions (*p = 0.0020*, 10.99 −150.02 Hz, −0.200 - 0.500 s from ripple center, corrected across time, frequency and voxels, masked by top 15% of voxels showing the strongest ripple-band effect). **B**, Coupling of MEG ripple-like activity to cortical SOs. Top, ripple occurrence relative to SO troughs, showing significantly increased ripple probability during the SO up-state (*p* = 0.0319, 0.20–0.65 s relative to SO troughs, corrected across time and sensors). Middle, topography of significant SO–ripple coupling sensors, maximal over fronto-temporal regions. Right, distribution of ripple events across the SO cycle, demonstrating significant alignment to the SO up-state (V-test against phase 0°, where 0 corresponds to the troughs in the SO-band: mean ± SEM = 17.82° ± 4.80°, *v* = 16.05, *p* < 0.001). **C**, Coupling of MEG ripple-like activity to cortical sleep spindles. Ripple occurrence was significantly modulated around spindle centers (most negative spindle trough), with maximal event density near the spindle center (*p* = 0.0339, −0.15 to 0.15 s relative to spindle center, corrected across time and sensors). Ripple events were significantly phase-locked to spindle troughs (V-test against phase ±π, where ±π corresponds to troughs in the spindle band: mean angle ± SEM = 176.12° ± 14.69°, *v* = 7.33, *p* = 0.007). **D**, Ripple band activity within SO-spindle complexes. Time–frequency analyses time-locked to the spindle trough of cortical SO–spindle complexes revealed the expected slow-oscillation and spindle activity. In the same analysis, MTL source-level activity showed a prominent ripple-band cluster centered around 90 Hz (80–120 Hz; 0–0.20 s from spindle center; *p* < 0.001, corrected across time, frequency, and voxels). Intracranial validation of MTL ripples. Grand-average ripple-locked activity and time–frequency representation computed from 38 MTL contacts across 15 patients. **F,** Coupling of intracranially recorded MTL ripples to cortical SOs and sleep spindles, which were detected from an additional 15 cortical contacts. MTL ripples occurred preferentially during the SO up-state (*p* = 0.01, 0.40–0.65 s from SO-trough) and were significantly aligned to SO the up-state phase (V-test vs phase 0: mean ± SEM = −5.93° ± 7.27°, *v* = 26.23, *p* < 0.001). MTL ripples were additionally coupled to spindle centers (*p* = 0.01, −0.15 to 0 s from spindle center, corrected across time) and significantly phase-locked to spindle troughs (V-test against phase ±π: mean ± SEM = 143.08° ± 10.96°, *v* = 9.26, *p* = 0.017).

We next examined whether source-reconstructed MEG ripple-like activity was embedded within the expected NREM oscillatory context defined by sensor-level cortical SOs and spindles (see Fig 2). To benchmark these dynamics against established physiological signatures of MTL ripples, we applied the same analytical approach to an independent intracranial EEG dataset comprising data from 15 patients, with intracranial MTL contacts (n = 38) used to identify ripples and cortical contacts (n = 15) used to detect SOs and sleep spindles (Fig. 3E). MEG ripple-like activity closely recapitulated the coupling dynamics observed for invasively recorded MTL ripples. Both MEG ripple-like activity and MTL ripples were significantly coupled to cortical SOs, with increased occurrence during the SO up-state (MEG: *p* = 0.032, 0.20 to 0.65 s relative to sensor-level SO trough, corrected across time and sensors, Fig. 3B, upper; iEEG: *p* = 0.01, 0.40 to 0.65 s relative to cortical SO trough, Fig. 3F, left). In

MEG, these SO-coupling effects were most prominent over fronto-temporal sensors (Fig. 3B, middle). Consistent with these peri-event analyses, ripple occurrence in both modalities was non-uniformly distributed across the SO cycle and significantly aligned to the SO up-state (V-test against phase 0; MEG: mean ± SEM = 17.82° ± 4.80°, v = 16.05, *p* < 0.001, Fig. 3B, right; iEEG: mean ± SEM = −5.93° ± 7.27°, v = 26.23, *p* < 0.001, Fig. 3F, bottom). A similar pattern was observed for spindle coupling. Both MEG ripple-like activity and MTL ripples were significantly coupled to spindle centers (most negative spindle trough), with maximal event density around the spindle center (MEG: *p* = 0.034, −0.15 to 0.15 s relative to sensor-level spindles , corrected across time and sensors, Fig. 3C/D; iEEG: *p* =0.032, −0.15 to 0 s relative to cortical spindles, corrected across time, Fig. 3F). In MEG, spindle-coupling effects were most prominent over temporo-parietal sensors. Ripple activity in both modalities also showed significant phase alignment to spindle troughs (V-test against phase ±π; MEG: mean angle ± SEM = 176.12° ± 14.69°, v = 7.33, *p* = 0.007, Fig. 3B lower; iEEG: mean ± SEM = 143.08° ± 10.96°, v = 9.26, *p* = 0.017, Fig. 3F right). Thus, source-reconstructed MEG ripple-like activity showed temporal and phase-coupling profiles that closely resembled those of invasively recorded MTL ripples, including preferential alignment to cortical SO up-states and spindle troughs.

Finally, we examined ripple-band activity in relation to complete SO-spindle complexes. We time-locked MEG source signals to the spindle center of sensor-level detected SO–spindle complexes and transformed the resulting epochs into the time–frequency domain. As expected, this analysis revealed elevated activity in the SO and spindle frequency ranges (1-50 Hz, −0.7 to 0.7 from spindle center, peaking at 13.5 Hz, *p* < 0.001, corrected across time, frequency, and voxels; Fig. 3D). Critically, it also identified a prominent ripple-band cluster centered around 94 Hz within MTL source voxels and time-locked to the spindle peak (50–130 Hz, 0 to 0.200 s from spindle center; *p* < 0.001, corrected across time, frequency, and voxels; Fig. 3D). Together, these findings show that MEG ripple-like activity is not only spatially concentrated in medial temporal regions, but also temporally aligned with spindles occurring during the SO up-state, placing it within the established SO–spindle–ripple hierarchy of NREM sleep.

### Targeted memory reactivation recruits MEG ripple-like activity alongside sleep spindles

Having established that MEG-derived ripple-like events show the spatial and temporal signatures expected of MTL ripples during spontaneous NREM sleep, we next asked whether they would be specifically engaged during TMR. TMR provides a controlled experimental manipulation of sleep-dependent memory processing, in which memory cues presented during NREM sleep reliably elicit cortical K-complexes (i.e. isolated SOs) and nested sleep spindles, with stronger cue-evoked spindle responses often observed for items that are later remembered^15,50,53,57,58^.

To test whether the MEG data captured the expected physiological response to memory cueing, we first compared cue-locked activity for items that were subsequently remembered versus not remembered after sleep. Time–frequency analyses of MEG sensor-level epochs revealed stronger spindle-band activity for subsequently remembered than not-remembered trials, with the effect centered in the canonical spindle range around 13 Hz and emerging between 0.9 and 2.5 s after cue onset (Fig. 4A; *p* = 0.04, cluster-corrected across time, frequency, and sensors). This enhancement emerged during the positive deflection of the cue-evoked K-complex (see white trace in Fig. 4A for a representative temporal sensor and Supplementary Fig. 3 for ERPs corresponding to remembered and not remembered cues), consistent with increased spindle recruitment during SO up-states. Event-based spindle analyses confirmed this pattern: spindle rates were significantly higher for remembered than not-remembered trials in a cluster ranging from 1.350 to 1.550 s after cue onset (*p* = 0.028, corrected across time and sensors; Fig. 4B; see Supplementary Table 2 for event rates). Thus, remembered trials were accompanied by the expected cortical spindle response ^15,50,59^, providing a physiological reference for subsequent source-level analyses.

**Figure 4.**
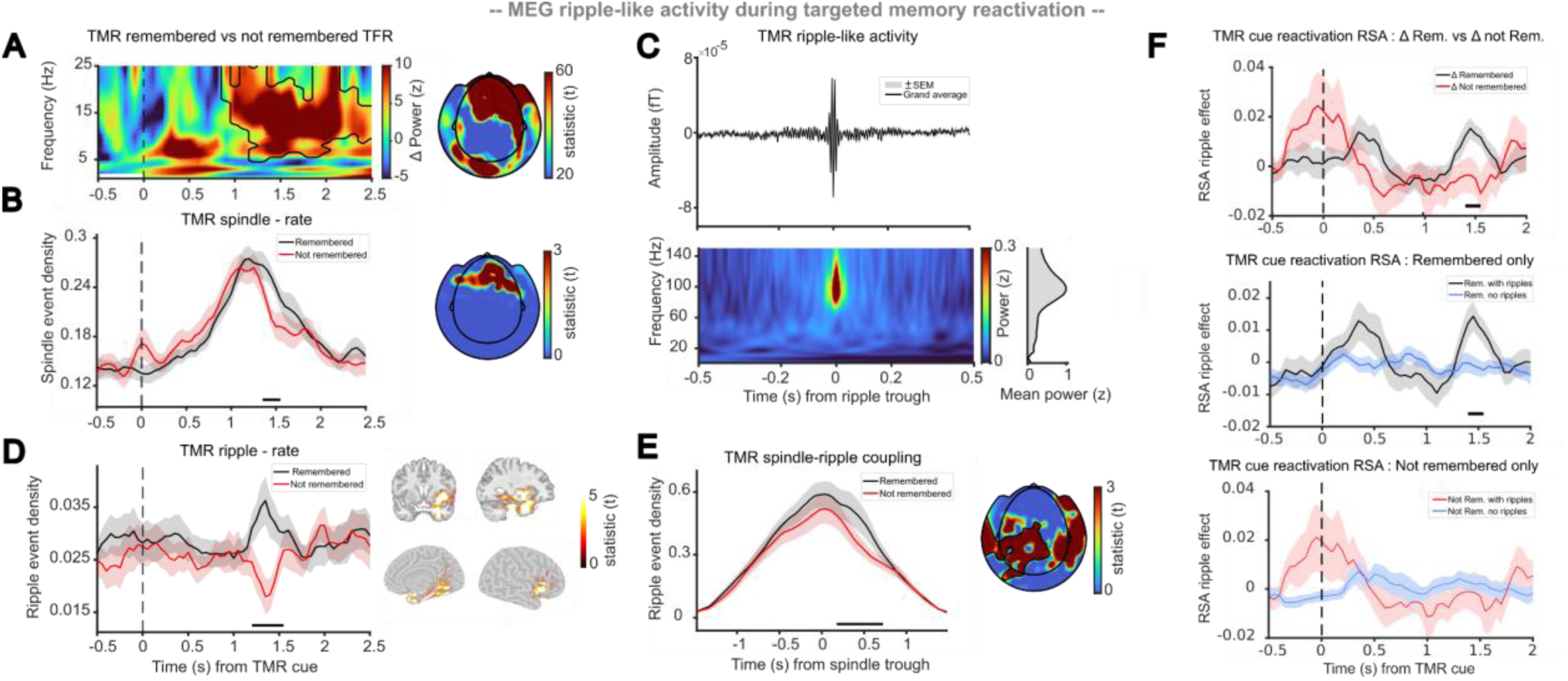
Targeted memory reactivation engages MEG ripple-like activity associated with subsequent memory retention. **A**, Cue-locked sensor-level TFR (remembered vs not-remembered items). TFR analysis revealed stronger activity for later remembered than not-remembered cues in a significant positive cluster spanning 0.90–2.50 s and 4.98–24.98 Hz relative to cue onset, peaking in the spindle range at 12.96 Hz and 1.50 s (cluster-corrected *p* = 0.04; corrected across time, frequency, and sensors). **B,** Cue-locked spindle-event analyses comparing subsequently remembered and not-remembered trials. Spindle rates were significantly higher for remembered trials in a cluster centered at 1.5 s after cue onset (*p*= 0.028, 1.350–1.550 s relative to cue onset, corrected across time and sensors). Data are shown as mean ± SEM.; see Supplementary Table 2 for event ratesaverage spindle density across sensors was 4.87 ± 0.26 events/30 s. **C,** Source-space detection of cue-evoked ripple-like activity. Grand-average ripple-like activity and corresponding time-frequency representation (see Supplementary Table 2 for event rates). . **D,** Memory-related modulation of MEG ripple-like activity during TMR. Ripple-like event rates were significantly higher for subsequently remembered than not-remembered items between 1.20 and 1.55 s after cue onset (*p* = 0.008, corrected across time and voxels). The effect was concentrated within medial temporal and downstream temporal regions. **E,** Coupling between TMR-related cortical spindle activity and source-level ripple-like activity. Remembered trials exhibited stronger spindle–ripple-like coupling than not-remembered trials just after the spindle peak (*p* = 0.049, 0.175–0.725 s, corrected across time and sensors), indicating enhanced coordination between cortical spindle activity and medial temporal ripple-like dynamics during successful memory reactivation. **F,** top panel, Ripple-associated item-specific reactivation in the medial temporal ripple-effect ROI (Fig. 4D). Reactivation was quantified as higher medial temporal pattern similarity for same-cue than different-cue trials. The plotted time courses show the ripple-related increase in same-cue RSA, defined as trials with ripple-like events minus trials without ripple-like events, separately for later remembered and not-remembered cues. The ripple-related RSA increase was significantly stronger for later remembered than not-remembered cues between 1.40 and 1.55 s after cue onset (memory outcome × ripple-status interaction: *p* = 0.044, corrected across time). The middle panel shows that this interaction was driven by later remembered cues, for which trials with ripple-like events exhibited higher same-cue RSA than trials without ripple-like events during the same interval (*p* = 0.048, corrected across time). No corresponding effect was observed for not-remembered cues (bottom panel; *p* = 0.263, corrected across time).

We next examined whether MEG ripple-like activity differed between subsequently remembered and not-remembered trials. Analogous to the spontaneous NREM analyses, ripple-like activity was detected voxel-wise in source space (Fig. 4C, see Supplementary Table 2 for event rates). Ripple-like event rates were significantly higher for remembered than not-remembered items between 1.20 and 1.55 s after cue onset (*p* = 0.008, corrected across time and voxels; Fig. 4D, see Supplementary Table 2 for event rates). This memory-related increase was predominantly localized to the MTL and downstream temporal regions (Fig. 4D; n = 71 significant voxels). Notably, it emerged in the same post-cue interval as the remembered–not-remembered difference in spindle activity, suggesting that both effects may reflect components of a shared cue-evoked sleep-oscillatory response^15^. We therefore tested whether cortical spindle activity and source-level ripple-like activity were more strongly coordinated during remembered as compared to not-remembered trials. Indeed, remembered trials showed stronger spindle–ripple-like coupling than not-remembered trials just after the spindle peak (*p* = 0.049, 0.175 to 0.725 s, corrected across time and sensors; Fig. 4E), indicating that memory cueing was associated not only with increased cortical spindle recruitment and enhanced MTL ripple-like activity, but potentially also with stronger coordination between these two signals.

Finally, we asked whether ripple-like events coincided with stronger TMR-induced representational specific reactivation. Because reminder cues were repeated six to seven times during TMR, we could compare activity patterns across repetitions of the same cue. Within the medial temporal ripple-effect ROI (Fig. 4D, n = 71 voxels), cue-specific reactivation was quantified as the difference between same-cue and different-cue pattern similarity, following the logic of representational similarity analysis^60–62^. We then tested whether this cue-specific RSA differed between trials with and without ripple-like events, separately for later remembered and not-remembered cues. This analysis revealed a significant memory outcome × ripple-presence (present vs. absent) interaction. Specifically, the ripple-related increase in cue-specific RSA was stronger for later remembered than for not-remembered cues between 1.40 and 1.55 s after cue onset (cluster-corrected across time; *p* = 0.044; Fig. 4F, top panel). Notably, this effect occurred within the same post-cue interval in which we observed memory-related increases in spindle and medial temporal ripple-like activity (Fig. 4 B and D), linking cue-specific reactivation to the oscillatory events identified in the preceding analyses. Follow-up analyses showed that this interaction was driven by remembered cues, for which trials with ripple-like events showed higher cue-specific RSA than trials without ripple-like events in the same time window (1.40–1.55 s, cluster-corrected across time; *p* = 0.048; Fig. 4F middle). No corresponding effect was observed for not-remembered cues (cluster-corrected across time; *p* = 0.263; Fig. 4F bottom).

Together, these results suggest that medial temporal ripple-like events marked moments of representational specific reactivation driven by the cue during successful TMR, rather than reflecting a general increase in source-pattern similarity.

## Discussion

We show that non-invasive MEG can track ripple-like medial temporal activity during human NREM sleep. This interpretation is supported by convergent evidence spanning (i) anatomical localization, (ii) embedding within the canonical slow oscillation–spindle hierarchy, (iii) correspondence with the physiological organization of intracranially recorded medial temporal ripples, and (iv) modulation during targeted memory reactivation. Together, these findings establish sleep MEG as a non-invasive approach for investigating ripple-related hippocampo-cortical dynamics across distributed brain networks in healthy humans.

Hippocampal ripples are central to sleep-dependent memory consolidation^12–16,45^, but in humans they have remained difficult to study outside intracranial recordings^12,27^. We therefore asked whether sleep MEG could provide a non-invasive window onto ripple-related medial temporal dynamics in healthy participants. Because transient ripple-band activity in source-reconstructed MEG is ambiguous, we did not define candidate events by spectral content alone, but instead evaluated their physiological plausibility against spatial, temporal, and mnemonic signatures derived from hippocampal ripple physiology^10,11,14–17^. Importantly, candidate events consistently fulfilled these independent physiological signatures: they were concentrated in medial temporal and downstream temporal regions (Fig. 3A), embedded within slow oscillation–spindle dynamics (Fig. 3B-D), closely resembled intracranially recorded MTL ripples (Fig. 3E, F), and were modulated by targeted memory reactivation (Fig. 4B-F). The convergence of these spatial, temporal, and mnemonic signatures therefore supports the interpretation that sleep MEG provides non-invasive access to physiologically meaningful ripple-related dynamics while extending their investigation beyond the spatially restricted sampling of intracranial recordings.

The findings build on growing evidence that medial temporal lobe activity is accessible to MEG. Evidence from simulation work and simultaneous iEEG–MEG recordings indicates that deep medial temporal signals can contribute to MEG recordings^31,32,40,63,64^. In parallel, MEG studies of episodic memory, working memory, language, and spatial navigation have reported hippocampal or broader medial temporal contributions to task-related oscillatory dynamics ^30,33,43,65–68^. The present study extends this work into sleep by showing that transient ripple-like events can be tracked within the physiological architecture of NREM sleep.

Invasive recordings in rodents and humans have been indispensable for discovering hippocampal ripples and defining their role in memory^12,20,27,45^. Across species, this work has shown that ripples contribute to the stabilization and redistribution of hippocampus-dependent memories and are temporally coordinated with cortical SOs and thalamo-cortical spindles as part of the canonical

NREM sleep hierarchy^7,8,10,11,14,20^. Within this hierarchy, ripple events are thought to provide temporally precise windows for hippocampo-cortical communication and memory reactivation during sleep^11,14–17^. Against this background, the present study asked whether key physiological signatures of ripple-related medial temporal dynamics can also be tracked non-invasively with MEG.

The first signature is spatial plausibility. Ripple-like MEG activity was not distributed randomly across source space, but was concentrated in medial temporal and downstream temporal regions. The extension beyond the medial temporal lobe should be interpreted cautiously, as it may partly reflect source leakage or the spatial spread inherent to MEG source reconstruction^69,70^. At the same time, previous electrophysiological, imaging, and intracranial recording studies have demonstrated that hippocampal ripples are associated with coordinated activity across distributed cortical networks, including temporal cortical regions ^4,7,71–75^. More direct evidence from human intracranial recordings indicates that ripple activity can be coordinated between the medial temporal lobe and temporal association cortex and is associated with successful memory retrieval and the reinstatement of task-specific cortical activity patterns during wake^76–78^. The spatial distribution of the present ripple-like events is also consistent with the known organization of hippocampal-neocortical communication pathways, whereby hippocampal output is relayed through the entorhinal, perirhinal, and parahippocampal cortices to distributed temporal and association neocortical regions^16,47,79,80^. Thus, their distribution across medial temporal and downstream temporal cortical regions in the present data is consistent with participation in a ripple-related hippocampal–cortical network rather than reflecting a nonspecific high-frequency signal.

The second signature is oscillatory embedding. Ripple-like events occurred preferentially during SO up-states and around spindle peaks, and source-level time–frequency analyses locked to sensor-detected SO–spindle complexes revealed a prominent ripple-band cluster in medial temporal voxels peaking during or shortly after the spindle peak (Fig. 3D). This temporal organization closely follows the canonical NREM sleep hierarchy established in invasive recordings in animals and humans^8–11^. Importantly, this correspondence was not only inferred from prior literature, but was supported by the present iEEG validation analysis: MEG ripple-like activity recapitulated key coupling dynamics of invasively recorded MTL ripples, including preferential occurrence during SO up-states and spindles. Thus, even without the local specificity of intracranial recordings, the alignment of ripple-like activity with SO–spindle complexes provides an important physiological constraint on the interpretation of the MEG signal. A natural next step will be to test this relationship experimentally. For example, closed-loop auditory stimulation could be used to enhance, shift, or perturb SO–spindle dynamics^81,82^ and assess whether ripple-like medial temporal activity is systematically modulated by these changes. Such an approach would help determine how closely the MEG signal follows experimentally induced changes in the sleep-oscillatory context.

The third signature is mnemonic relevance. In the targeted memory reactivation analysis, ripple-like MEG activity was not only embedded in NREM sleep oscillatory dynamics, but was preferentially enhanced for cues that were later remembered. In line with previous literature, remembered cues elicited stronger spindle-band activity and higher spindle event rates than not-remembered cues, with event-based spindle effects peaking around 1.35–1.55 s after cue onset^53^. Ripple-like event rates showed a parallel memory-related increase between 1.20 and 1.55 s after cue onset, predominantly in medial temporal and downstream temporal regions. Critically, spindle-triggered analyses further revealed stronger spindle–ripple-like coupling for remembered than not-remembered trials. Thus, successful cueing was associated not simply with stronger cortical spindle recruitment or stronger medial temporal high-frequency activity in isolation, but with enhanced coordination between the two. The timing of this effect aligns with human work showing that memory reactivation is preferentially detected during SO–spindle complexes^16,52^, particularly in the post-downstate interval when spindles are nested in the excitable phase of the SO^55^.

The RSA analysis adds an important functional constraint to this interpretation. Rather than showing only that ripple-like events were more frequent during successful cueing, this analysis linked them to cue-specific pattern similarity within the same medial temporal ROI. Trials containing ripple-like events showed a stronger same-cue similarity increase for later remembered than for not-remembered cues, suggesting that the detected events were associated with the content-specific component of TMR-related processing. This strengthens the interpretation that MEG ripple-like activity reflects physiologically meaningful medial temporal dynamics, rather than nonspecific high-frequency activity or a generic response to auditory stimulation. In line with intracranial evidence that spindle-locked MTL ripples carry information about reactivated memories during NREM sleep^15,16^, the present findings suggest that sleep MEG can capture a non-invasive signature of coordinated spindle– ripple-like windows in which memory-related patterns are preferentially reinstated.

More broadly, our findings position sleep MEG as a translational bridge between invasive ripple physiology and scalable human cognitive neuroscience. Scalp EEG has been essential for linking SO– spindle coupling to memory consolidation in healthy participants, but provides limited access to fast medial temporal activity. Intracranial recordings provide local specificity, but are restricted to clinical populations and sparse electrode coverage. MEG occupies an intermediate position: it cannot provide the cellular or local field specificity of iEEG, but it can be applied non-invasively while preserving millisecond-scale temporal resolution and offering source-level sensitivity to medial temporal dynamics. This opens a route for studying ripple-like hippocampo-cortical coordination across healthy populations, aging cohorts, and clinical or interventional settings where invasive recordings are not feasible.

## METHODS

### Participants

Eighteen healthy, right-handed native Dutch speakers participated in the study (age: 23.94 ± 0.97 years, mean ± SEM; 9 female). Five additional participants were excluded because they did not achieve stable NREM sleep during MEG recordings (3 female). Participants were free of medication and reported no history of neurological or psychiatric disorders. All participants reported good sleep quality and regular sleep-wake cycles. Prior to each experimental session, participants were instructed to restrict sleep to 6 hours, avoid caffeine and alcohol, and refrain from daytime naps. The study was approved by the local ethics committee (CMO-2014/288, region Arnhem-Nijmegen), and all participants provided written informed consent before participation. Participants received monetary compensation after completing the experiment. Eighteen participants completed the full protocol. However, due to sleep-related difficulties during the experimental TMR night that prevented reliable cue delivery, TMR analyses were restricted to 16 participants. In addition, a complementary intracranial EEG (iEEG) dataset from 15 patients with medically intractable epilepsy (8 female; age: 30.33 ± 2.36 years, mean ± SEM) was used as a complementary benchmark dataset for the canonical coupling profile of MTL ripples during NREM sleep. Patients were recruited at the Epilepsy Center, Department of Neurology, Ludwig-Maximilians-Universität München (Munich, Germany). The iEEG study was approved by the ethics committee of the Medical Faculty of Ludwig-Maximilians-Universität München. All patients provided written informed consent.

### Experimental Design and Procedure

The experiment comprised three sessions per participant. First, participants completed an adaptation session lasting approximately 1 hour, which served to habituate them to the experimental setup and sleeping in the MEG environment. The adaptation session took place approximately 1 week before the first experimental session. Participants then completed two experimental sessions, each comprising a pre-sleep memory task, an initial recall test (Recall Pre), a sleep interval of approximately 90 minutes, and a post-sleep recall test (Recall Post). The two experimental sessions were separated by approximately 1 week, followed the same task structure but used different item sets, and differed only in whether targeted memory reactivation (TMR) was applied during sleep. In one session, auditory memory cues associated with a subset of learned items were presented during NREM sleep (TMR session); in the other, no cues were presented (non-TMR session). The order of TMR and non-TMR sessions was counterbalanced across participants. In both experimental sessions, EEG and EOG electrodes were applied at 20:30 h. Around 21:30 h, participants entered the MEG system and completed the memory task, followed by an initial recall test (Recall Pre). The sleep interval started at approximately 22:30 h. Participants were awakened from NREM sleep after approximately 90 minutes of sleep, followed by a final recall test (Recall Post).

*Memory Task.* In each experimental session, participants learned a different set of 140 Dutch word-picture associations. Each Dutch word was paired with a neutral object image. All images were taken from ^83^. Learning consisted of two rounds. During the encoding round, each trial began with a fixation cross accompanied by auditory presentation of the word cue (1500 ms), followed by brief presentation of the associated image (300 ms) on either the left or right side of the screen (randomized). This was followed by a jittered fixation interval (2800-3000 ms). After 70 trials, participants were offered a short break. In the second learning round, participants completed a cued recall task with feedback. Each trial began with a jittered fixation cross (2300-2700 ms), followed by auditory presentation of the word cue (2000 ms). A question mark then prompted verbal recall. After the response, the correct image was briefly presented as feedback (350 ms) on the corresponding side of the screen. Memory performance was assessed using cued recall without feedback. The procedure was identical to the cued recall task during learning, except that no feedback image was presented after the response. An initial recall test (Recall Pre) was administered after the memory task and before sleep. A final recall test (Recall Post) was administered after the sleep interval.

*Targeted Memory Reactivation (TMR).* During the TMR session, 70 of the 140 learned words were presented again during the retention interval. Cue selection was based on pre-sleep recall (Recall Pre) performance, such that the proportions of remembered and non-remembered items were matched between the cued and uncued conditions. Words were presented binaurally at approximately 50 dB SPL via ear inserts. Cues occurred every 5800–6200 ms for approximately 60 minutes, resulting in 6– 7 repetitions per cue. During sleep, cues were delivered during NREM sleep (N2 and slow-wave sleep). Stimulation was paused whenever polysomnography indicated arousal, wakefulness, or REM sleep.

*Sleep EEG.* EEG was recorded simultaneously with MEG using Ag/AgCl electrodes placed at F3, F4, C3, C4, and A2, referenced to A1 (left mastoid). Horizontal and vertical EOG were recorded bipolarly. Signals were sampled at 600 Hz, and impedances were kept below 20 kΩ. Sleep stages were monitored online and scored offline by two independent raters according to standard criteria^22^. Offline scoring used average mastoid referencing and band-pass filtering from 0.1 to 30 Hz.

*MEG Recordings.* MEG data were acquired at the Donders Institute for Brain, Cognition and Behaviour, Radboud University, Nijmegen, the Netherlands. Brain activity was recorded using a whole-head 275-channel MEG system (VSM/CTF) in a magnetically shielded room. Data were sampled at 600 Hz. Head position was tracked continuously using localization coils placed at the nasion and bilateral ear canals. Whenever head displacement exceeded 5 mm, a new recording file was initiated to ensure stable head position within individual recording segments. For source reconstruction, high-resolution structural MRI scans were acquired for each participant using a 3T Siemens scanner.

*iEEG Recordings.* Intracranial EEG was recorded from Spencer depth electrodes (Ad-Tech Medical Instrument, Racine, Wisconsin, United States) with 4–12 contacts each, 5 mm apart. Data were recorded using XLTEK Neuroworks software (Natus Medical, San Carlos, California, US) and an XLTEK EMU128FS amplifier, with voltages referenced to a parietal electrode site. The sampling rate was set at 1000 Hz.

### Data Analysis

All data were analyzed using Matlab (2018b and 2024a; Mathworks). MEG and iEEG data were preprocessed using the FieldTrip toolbox^84^, v.09/01/2020 and v.15/10/2023). For both MEG and iEEG data, artefactual segments were identified by visual inspection and excluded from further analyses. For the MEG dataset, SQUID jump artefacts were additionally detected automatically and rejected prior to event detection. For intracranial analyses, contacts in the medial temporal lobe (MTL; comprising hippocampus, parahippocampal cortex, and entorhinal cortex) were selected based on anatomical localization, yielding 2.53 ± 0.48 contacts per patient across the 15 patients (mean ± SEM). In addition, one cortical reference contact per patient was selected as the channel exhibiting the highest sleep spindle power. For the iEEG dataset, interictal epileptiform discharges (IEDs) were then detected separately for each channel (25–80 Hz Hilbert envelope; z > 3; 20–100 ms), and channel-specific ±0.5-s intervals around detected IEDs were excluded from all subsequent analyses^8,85,86^.

*Behavior.* In each experimental session, memory performance was assessed for 140 word–picture associations. As an index of memory retention across the sleep interval, we calculated the relative change in the number of correctly recalled word–picture associations from before to after sleep, with pre-sleep recall performance (Recall Pre) set to 100%. For the non-TMR session, this measure was computed across all 140 associations. For the TMR session, retention was additionally computed separately for cued and uncued items.

*Source-level analyses.* To reconstruct source-level time series, we used a virtual electrode approach based on linearly constrained minimum variance (LCMV) beamforming^56^, as implemented in FieldTrip. Individual structural MR images were aligned to the MEG coordinate system based on participant-specific head shapes. A realistic single-shell brain model^87^ was constructed for each participant from the individual structural MRI. Source estimates were interpolated onto the individual anatomical images. The LCMV beamformer, which computes a spatial filter based on the lead field of each source location and the covariance matrix of the sensor-level data, was used to reconstruct source time series. For the non-TMR session, all artifact-free NREM sleep segments were projected into source space to characterize spontaneous sleep activity. For the TMR session, sensor-level data were projected into source space in cue-locked epochs from −2 to +5 s relative to auditory cue onset (481.94 ± 24.61 trials per participant, mean ± SEM) to assess cue-related responses during sleep. Applying the spatial filters to the sensor-level data yielded source time series for 1,457 virtual sensors covering the brain volume.

*Event detection.* SOs, sleep spindles, and ripples were detected separately in the MEG and iEEG datasets for each participant and condition using established algorithms^11,25,55^. For the iEEG dataset, event detection was restricted to artifact-free NREM sleep (N2 and SWS). SOs and sleep spindles were detected on the cortical reference contact, whereas ripples were detected on all MTL contacts. For SO detection, signals were band-pass filtered between 0.3 and 1.25 Hz, and candidate events with durations between 0.8 and 2 s were identified. Of these, only events exceeding an amplitude threshold corresponding to the mean amplitude of detected events plus 1.5 SD entered the analysis. For spindle detection, signals were band-pass filtered between 12 and 16 Hz. An RMS envelope (200-ms window) was computed, and candidate events with durations between 0.5 and 3 s were identified. Of these, only events whose RMS amplitude exceeded the median amplitude of detected events by a factor of 1.3 entered the analysis. For ripple detection, signals were notch filtered around 50, 100, and 150 Hz (±1 Hz) and band-pass filtered between 80 and 120 Hz. An RMS envelope (20-ms window) was computed, and candidate events with durations between 38 and 500 ms were identified. Of these, only events exceeding an RMS threshold corresponding to the mean amplitude of detected events plus 2.5 SD entered the analysis.

For the MEG dataset, SOs and sleep spindles were detected at the sensor level using the same detection parameters as for the iEEG cortical reference contact. In the non-TMR condition, detection was performed on continuous artifact-free NREM data, whereas in the TMR condition, detection was performed on appended cue-locked trials spanning −2 to +5 s relative to auditory cue onset. Ripple-like activity was detected in source space using the same spectral and temporal detection parameters as for iEEG ripples, based on the corresponding continuous (non-TMR) or cue-locked (TMR) source-space data described above.

*Source-space ripple segmentation and time–frequency analysis (non-TMR condition)*. Ripple-like events detected independently in each of the 1,457 source voxels were consolidated into a common set of ripple-associated episodes across source space to preserve the interpretability of spatial power differences. Voxel-wise ripple detections were merged using a neighbourhood-constrained spatial-consistency procedure based on a precomputed source-space adjacency structure. Specifically, detections were evaluated within non-overlapping 5-s windows, and a window was retained only when ripple-like events were present in at least 50% of neighbouring source voxels within a 2-cm radius. To avoid counting spatially extended detections multiple times, each accepted window contributed a single representative ripple-like event time, sampled from the event minima detected across the contributing source voxels. Events occurring within a 2-s refractory period were removed. For each retained episode, an 8-s epoch (−4 to +4 s) was extracted from all source voxels.

Source-space data were detrended, demeaned, high-pass filtered at 0.5 Hz, and notch filtered at 50, 100, and 150 Hz (±2 Hz). Time–frequency representations were computed using a Hanning-tapered sliding-window approach with 4 cycles per frequency, from 1–30 Hz in 1-Hz steps and 35–150 Hz in 5-Hz steps, and from −2 to +2 s in 100-ms steps. Single-trial power was z-scored separately for each participant, voxel, and frequency across all trials and the interval from −2 to +2 s, and then averaged across trials. Ripple-associated estimates were contrasted against surrogate non-events matched in number and sleep stage (N2/SWS), sampled from non-ripple periods within a 300-s window surrounding each ripple event, and processed identically. Group-level differences between ripple-associated and surrogate TFRs were assessed using two-sided cluster-based permutation testing as implemented in FieldTrip ^84,88^, based on a dependent-samples t-statistic, across the interval from −1 to +1 s relative to ripple peak and the frequency range from 1 to 150 Hz (500 permutations; cluster alpha = 0.05). Following this event-locked source-space TFR contrast, voxels were ranked according to their ripple-band statistical effect in the 80–120 Hz range, and the 15% of voxels showing the strongest ripple-band power increase relative to surrogate events were retained as a data-driven hotspot mask for subsequent non-TMR analyses (n = 219; Fig. 3A).

*Cue-locked time–frequency analysis (TMR condition).* For the TMR condition, cue-locked time– frequency representations were computed at the sensor level to characterize oscillatory responses to auditory memory cues during sleep. Analyses were based on artifact-free cue-locked trials. Time– frequency representations were computed across all MEG sensors using a Hanning-tapered sliding-window approach with 4 cycles per frequency, from 1 to 40 Hz in 1-Hz steps and from −2 to +4 s in 100-ms steps. Power estimates were then averaged separately for subsequently remembered and not-remembered cues, and baseline-corrected relative to the pre-cue interval (−0.5 to 0 s). Group-level differences between remembered and not-remembered cues were assessed using two-sided cluster-based permutation testing across sensors, frequencies (1–25 Hz), and time (−0.5 to +2.5 s), using a dependent-samples t-statistic (500 permutations, cluster alpha = 0.05).

*Peri-event time histograms.* To quantify the temporal relationship between slow oscillations (SOs), sleep spindles, and ripples/ripple-like events, peri-event time histograms (PETHs) were computed for both the iEEG and MEG datasets. For event-triggered analyses, SO timing was defined by the SO downstate, whereas spindle and ripple timing were defined by the sample of minimum amplitude. For the iEEG dataset, PETHs were computed for SO–spindle, SO–ripple, and spindle–ripple coupling within a ±1.5-s window using 50-ms bins. SOs and spindles detected on the cortical reference contact served as trigger events, and spindle or ripple occurrence was quantified relative to these triggers. For SO– spindle analyses, spindle occurrence was quantified on the same cortical reference contact. For SO– ripple and spindle–ripple analyses, ripple occurrence was quantified separately for each available MTL contact and then averaged across MTL contacts to yield a single cortical–MTL coupling profile per cortical reference contact.

For the MEG non-TMR condition, PETHs were likewise computed for SO–spindle, SO–ripple-like, and spindle–ripple-like coupling within a ±1.5-s window using 50-ms bins. Sensor-level SOs and spindles served as trigger events. For SO–spindle analyses, spindle occurrence was quantified at the sensor level, whereas for SO–ripple-like and spindle–ripple-like analyses, ripple-like activity was quantified in source space across the 15% strongest hotspot voxels (219 voxels), defined based on the event-related ripple power increase identified in the source-space ripple segmentation and time– frequency analysis (Fig. 3A).

For the MEG TMR condition, cue-locked PETHs were computed separately for subsequently remembered and not-remembered cues to quantify SO, spindle, and ripple-like event occurrence relative to auditory cue onset. SOs and spindles were quantified at the sensor level, whereas ripple-like activity was quantified in source space across all 1,457 voxels. In addition, spindle–ripple-like coupling during TMR was assessed by computing spindle-triggered PETHs separately for subsequently remembered and not-remembered trials, using sensor-level spindles as trigger events and ripple-like events from the significant source-voxel set as target events. For this analysis, both spindle and ripple-like events were restricted to the post-cue interval from 0.5 to 2.0 s, and PETHs were computed within a ±1.5-s window using 50-ms bins.

Statistical significance was assessed using two-sided cluster-based permutation testing (500 permutations; cluster alpha = 0.05). For the iEEG benchmark analyses, contact-level statistics were used to characterize canonical SO–ripple and spindle–ripple coupling profiles of invasively recorded MTL ripples. For iEEG and MEG non-TMR PETHs, event rates were tested against a null model defined by the time-averaged event rate of each individual PETH. For TMR analyses, subsequently remembered and not-remembered cues were compared directly. Correction was performed across time for iEEG PETHs, across time and sensors for MEG sensor-level analyses, and across time and voxels for MEG source-level ripple-like analyses. For TMR spindle–ripple-like coupling, correction was performed across time and sensors.

*Preferred phase analyses.* To further characterize the temporal embedding of spindle and ripple events within ongoing slow oscillatory activity, preferred-phase analyses were performed in the iEEG dataset and in the non-TMR MEG dataset using Hilbert-based phase estimates. In the iEEG dataset, instantaneous SO phase was extracted from continuous NREM data after band-pass filtering between 0.3 and 1.25 Hz. Analyses were restricted to periods within ±1.25 s of SO down-states defined from the cortical SO channel. SO phase values were then sampled at ripple times from each MTL contact and at spindle times from the analyzed contacts. In the non-TMR MEG dataset, preferred-phase analyses were likewise based on continuous artifact-free NREM data. For SO–spindle analyses, SO phase (0.3–1.25 Hz) was extracted from sensor-level data and evaluated within sensor-specific windows of ±1.25 s around detected SO down-states, and phase values were sampled at spindle times on the same sensor. For SO–ripple analyses, SO phase was extracted in the same frequency range but restricted to a common ±1.25-s window around SO down-states from the reference SO channel; phase values were then sampled at ripple event times from the previously defined source-space ripple voxel set used in the non-TMR MEG PETH analyses. For spindle–ripple analyses, spindle-band phase (12–16 Hz) was extracted from sensor-level data and restricted to windows of ±0.5 s around detected spindle centers; phase values were then sampled at ripple event times from the same source-space ripple voxel set. Phase angles were defined such that 0° corresponded to the SO up-state, whereas ±π rad (180°) corresponded to the spindle trough, defined as the most negative spindle deflection. At the group level, statistical significance was assessed using the circular V-test against 0° for SO–spindle and SO–ripple analyses, corresponding to preferential alignment with the SO up-state^8^. For spindle–ripple analyses in the non-TMR MEG dataset, the circular V-test was performed against ±π rad (180°), corresponding to preferential alignment with the spindle trough^8^.

*Source-level representational similarity analysis.* To test whether ripple-like events were associated with cue-specific memory reactivation, we performed a source-level representational similarity analysis (RSA) within the medial temporal ripple-effect ROI identified in the cue-locked ripple-event analysis (71 source voxels). For each participant, source activity was restricted to these voxels. Activity patterns were extracted in sliding temporal windows centered from −0.5 to 2.5 s relative to cue onset, using a 0.30 s window and a 0.05 s step size. Each pattern consisted of activity across ROI voxels and samples within the local time window.

For each time point, trial-wise activity patterns were normalized across all voxel-by-time samples within the local window and correlated across trials. Correlation coefficients were Fisher-z transformed. Cue-specific reactivation was quantified as the difference between the average similarity of trials belonging to the same reminder cue and the average similarity of trials belonging to different reminder cues. Only cues with at least two repetitions were included. RSA time courses were computed separately for later remembered and not-remembered cues. To test whether cue-specific reactivation differed between trials with and without ripple-like events, trials were classified as ripple-containing when at least one ROI voxel contained a detected ripple-like event in the post-cue window from 0.8 to 1.5 s after cue onset. This window encompassed the post-cue interval in which cue-locked ripple-like activity was observed in the preceding analysis (Fig. 4D). Same-cue RSA was computed separately for trials with and without ripple-like events, within later remembered and not-remembered cues. The ripple-related RSA increase was defined as the difference between these two estimates. The primary statistical test assessed whether this increase differed between later remembered and not-remembered cues, corresponding to a memory outcome × ripple-status interaction. Statistical inference was performed across participants using cluster-based permutation testing over time. Paired-sample t-tests were computed at each time point, and cluster-level significance was assessed using Monte Carlo permutations. The same procedure was used for follow-up tests comparing trials with and without ripple-like events separately for later remembered and not-remembered cues (500 permutations; cluster alpha = 0.05).

## Acknowledgements

T. Sc. is supported by the Emmy Noether program of the German Research Foundation (492835154). F.S. is supported by the South Tyrolean Fund for the Promotion of scientific Research (SFPR) at the South Tyrolean Health Care Service (SABES) and the Paracelsus Medical University Salzburg (PMU) (CUP I33C24008120003). C.F.D. is supported by the Max Planck Society. MEG data were acquired at the Donders Centre for Cognitive Neuroimaging, Donders Institute for Brain, Cognition and Behaviour, Radboud University, Nijmegen, the Netherlands.

## Data availability

All data supporting the findings of this study will be made publicly available upon publication.

## Code availability

All code related to the analyses of the manuscript will be made publicly available upon publication.

## Author contributions

T.Sc., O.J., and C.F.D. conceptualized and designed the study. T.Sc. acquired the data. J.R. and C.V. provided the clinical iEEG data. F.S. and T.Sc. analyzed the data. F.S. and T.Sc. interpreted the findings and wrote the original draft. All authors reviewed, edited, and approved the final manuscript.

