## Supplemental Information for "Non-invasive tracking of ripple-like activity during human sleep using MEG"

### Supplementary Information

| | noTMR (n = 18) | TMR (n = 16) | iEEG (n = 15) |
|---|---|---|---|
| **Wake (%)** | 4.68 ± 2.04 | 5.95 ± 3.52 | 7.65 ± 2.13 |
| **N1 (%)** | 11.19 ± 2.36 | 6.13 ± 1.56 | 4.00 ± 0.63 |
| **N2 (%)** | 54.38 ± 3.71 | 59.63 ± 4.48 | 47.79 ± 2.09 |
| **N3 (SWS, %)** | 27.40 ± 4.64 | 24.66 ± 3.75 | 18.87 ± 2.00 |
| **REM (%)** | 1.42 ± 0.66 | 2.87 ± 1.29 | 19.19 ± 1.78 |
| **Total sleep time (min)** | 93.03 ± 4.98 | 105.59 ± 5.32 | 477.20 ± 15.68 |

**Supplementary Table 1 | Sleep architecture across experimental conditions or cohorts** *(mean ± SEM)***.** Values are presented as *mean ± standard error of the mean (SEM)*. n indicates the number of participants contributing data per cohort. SWS = slow-wave sleep. REM = rapid eye movement sleep.

| Event Type | Condition | Rate (Mean ± SEM) | Unit |
|---|---|---|---|
| **Spindle** | All trials | 0.6780 ± 0.0239 | events/sensor/trial |
| | Remembered | 0.6785 ± 0.0244 | events/sensor/trial |
| | Not remembered | 0.6741 ± 0.0245 | events/sensor/trial |
| **Ripple-like** all voxels | All trials | 0.1041 ± 0.0094 | events/voxel/trial |
| | Remembered | 0.1072 ± 0.0119 | events/voxel/trial |
| | Not remembered | 0.1013 ± 0.0082 | events/voxel/trial |
| **Ripple-like** hotspot voxels | All trials | 0.1104 ± 0.0109 | events/voxel/trial |
| | Remembered | 0.1128 ± 0.0119 | events/voxel/trial |
| | Not remembered | 0.1051 ± 0.0110 | events/voxel/trial |

**Supplementary Table 2 | Descriptive statistics for spindle and ripple-like events during TMR.** Values are reported as mean ± SEM across participants. Event rates are provided for all trials and separately for subsequently remembered and not-remembered trials. For ripple-like events, rates are reported for all voxels (n = 1457) and for the subset of hotspot voxels showing significant differences in event rates between remembered and not-remembered trials (Fig. 4D; n = 71).

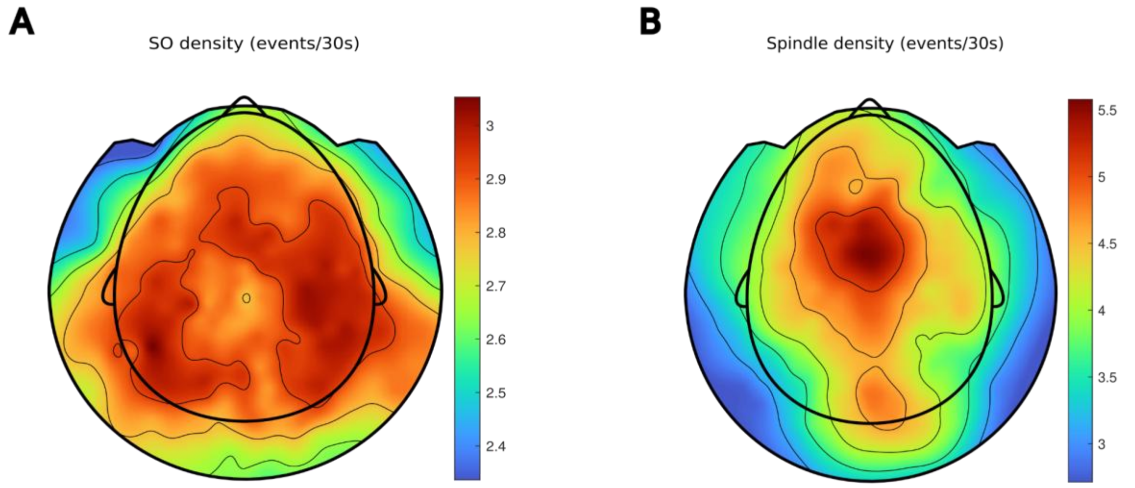


**Supplementary Fig. 1 | Topographical distributions of sleep event densities. A,** Sensor-level SO density and **B,** sensor-level spindle density averaged across participants during a nighttime nap measured with MEG.

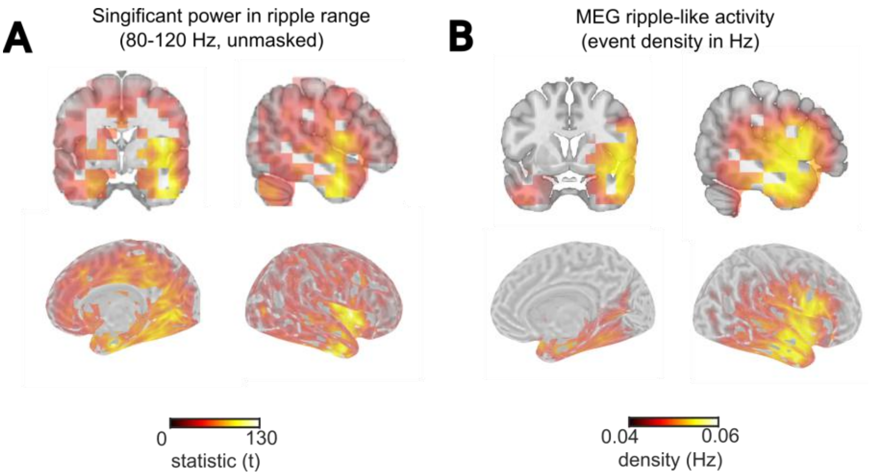


**Supplementary Fig. 2 | Convergent spatial localization of NREM ripple-like activity. A,** Voxel-wise statistical map of ripple-band (80–120 Hz) power associated with ripple-like episodes relative to surrogate events. Significant voxels with highest effects were centred on the MTL and downstream temporal cortical regions ($p$ = 0.002, 10.99 to 150.02 Hz, -0.200 to 0.500 s from ripple center, corrected across time, frequency and voxels. **B,** Voxel-wise density map of ripple-like events detected during NREM sleep. Regions with the highest event density overlapped closely with the power-based localization shown in (A), with maxima in the MTL and downstream temporal cortex.

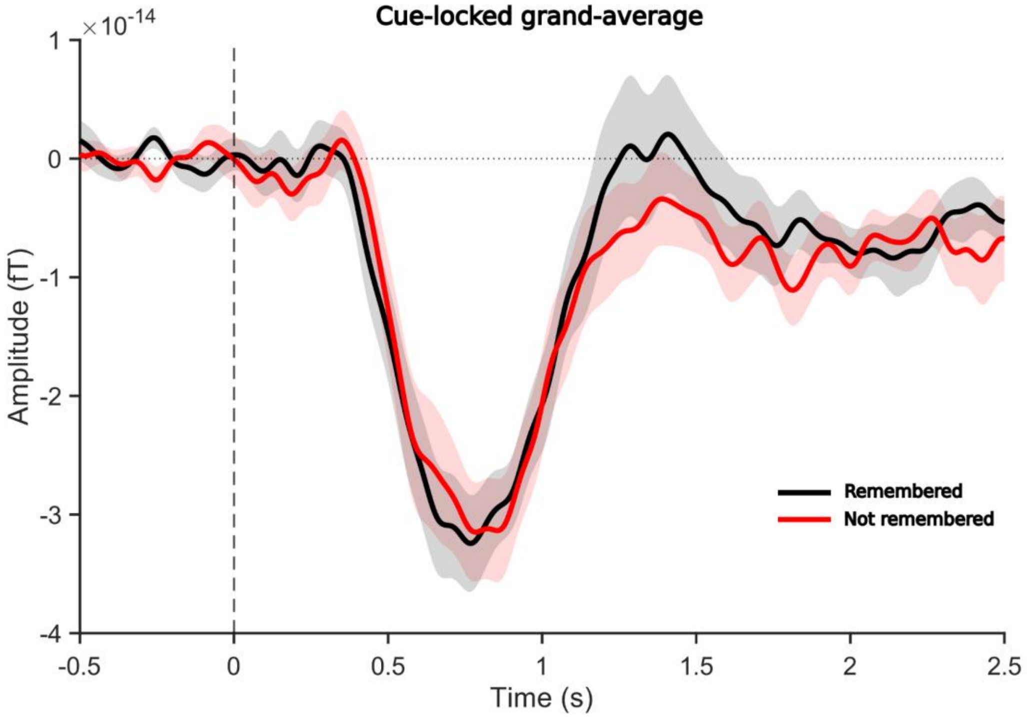


**Supplementary Fig. 3 | TMR cue-locked grand-average neural response on sensor level.** Sensor neural signals time-locked to TMR cues averaged for subsequently remembered (black) and not remembered (red) trials. Both conditions exhibit the characteristic K-complex response evoked by TMR cue presentation during NREM sleep.
